# The orphan receptor IL-17RD is a negative regulator of RIG-I-like receptor-dependent antiviral innate immunity and restrains SARS-CoV-2-induced lung inflammation

**DOI:** 10.64898/2026.09.01.747870

**Authors:** Firas El-Mortada, Jiunn Roy, Charlotte Girondel, Florence Dô, Ana Cláudia Dos Santos Pereira Andrade, Emile Lacasse, Isabelle Dubuc, Philippe Roux, Guillaume St-Jean, Baptiste Demey, Nathalie Labrecque, Louis Flamand, Sylvain Meloche, Marc J. Servant

## Abstract

Detection of viral RNA by the RIG-I-like receptors (RLRs) RIG-I and MDA5 triggers assembly of a MAVS-dependent signalosome that activates the TBK1-IRF3 and IKKβ-NF-κB axes together with the JNK and p38 MAPK modules, driving type I and type III interferon (IFN) and inflammatory cytokine production. Because unrestrained activity of this pathway is a major cause of immunopathology, host-encoded negative regulators are essential, yet the full complement of these brakes remains incompletely defined. Here we identify interleukin-17 receptor D (IL-17RD, also known as SEF), an orphan member of the IL-17 receptor family previously characterized as an antagonist of FGF and Toll-like receptor signaling, as a negative regulator of RLR-driven antiviral innate immunity. Using a CRISPR-engineered and shRNA-depleted human airway epithelial-derived lung carcinoma A549 cell line, we show that loss of IL-17RD amplifies and prolongs phosphorylation of TBK1 and IRF3 in response to poly I:C transfection and to infection with encephalomyocarditis virus (EMCV) or Sendai virus (SeV), and likewise potentiates the IKKβ-IκBα module and the TAK1-JNK1/2 and p38 MAPK branches. This translates into increased nuclear accumulation of IRF3 and p65, and markedly elevated induction of *IFNB1*, *IFNL1-3*, *CCL5*, *IL6*, and *NFKBIA* transcripts, as well as secreted IFN-β and IL-6. Silencing IL-17RD in ACE2-expressing A549 cells similarly derepresses the antiviral and inflammatory transcriptional programme following SARS-CoV-2 infection. Epistasis experiments place IL-17RD at the level of MAVS, downstream of the RLR sentinels. Mechanistically, IL-17RD localizes to the ER-to-Golgi intermediate compartment (ERGIC), the membrane platform on which the MAVS signalosome is present, and associates with RIG-I, MDA5, MAVS, TBK1 and IRF3. Its re-expression in depleted cells redistributes RLR effectors and TRAF proteins across low-molecular-weight signalosome fractions, reducing the amount of IRF3 recruited to the 670 kDa MAVS signalosome complex. Complementation of IL-17RD-deficient cells also indicates that the intracellular TIR subdomain is sufficient to confer this antagonistic activity. Finally, *Il17rd*^−/-^ mice display a splenic transcriptome enriched for antiviral response signatures, and, following intranasal infection with a moderate dose of SARS-CoV-2, they mount an exaggerated pulmonary cytokine response and develop significantly greater lung inflammation and fibrosis than wild-type littermates. Together, these data establish IL-17RD as a bona fide brake on the RLR-MAVS axis that limits virus-induced immunopathology, and identify the SEFIR/TIR subdomain as the module responsible for this activity.

## INTRODUCTION

The innate immune system provides the first line of defence against viral pathogens. Cytosolic recognition of viral RNA is mediated primarily by the RIG-I-like receptors (RLRs), a family of DExD/H-box helicases comprising RIG-I (DDX58), MDA5 (IFIH1) and LGP2 (DHX58) (1–3). RIG-I preferentially binds short double-stranded RNA with a 5′-triphosphate, as generated during the replication of negative-strand RNA viruses such as Sendai virus (SeV). By contrast, MDA5 senses long double-stranded RNA, characteristic of picornaviruses such as encephalomyocarditis virus (EMCV) and of the synthetic mimetic polyinosinic:polycytidylic acid (poly I:C). Ligand binding relieves autoinhibition, promotes K63-linked polyubiquitination and oligomerization of the receptors, and enables interaction with the mitochondrial antiviral signaling protein MAVS. MAVS serves as the nucleating platform for a higher-order signaling complex. Contrary to its original description as a strictly mitochondrial adaptor, functional MAVS signalosomes also assemble near or at other biological membrane structures, such as the ER-to-Golgi intermediate compartment (ERGIC) and peroxisomes (4, 5). Within this complex, TRAF2, TRAF3, TRAF5 and TRAF6 serve as ubiquitin-dependent scaffolds that recruit and activate two kinase modules: the IKK-related kinases TBK1 and IKKε, which phosphorylate IRF3 on its C-terminal serine cluster, driving dimerization, nuclear accumulation and transcription of IFNB1 and interferon-stimulated genes (6–8), and the IKK complex, which triggers IκBα degradation and NF-κB-dependent inflammatory gene expression (9). The intervening kinase architecture of the MAPK arm is, however, stimulus- and cell-type-dependent rather than a single canonical relay (10–13). The coordinated action of these branches produces type I and type III IFNs, chemokines such as CCL5, and pro-inflammatory cytokines including IL-6.

The magnitude and duration of this response must be tightly controlled. Insufficient IFN production permits uncontrolled viral replication, as illustrated by severe COVID-19 phenotypes associated with inborn errors of type I IFN immunity or with neutralizing anti-IFN autoantibodies (14, 15). Conversely, excessive or prolonged signaling drives tissue damage and hyperinflammation. Rather than viral burden per se, hyperinflammation is a principal determinant of mortality in severe respiratory viral disease (16–20). A large repertoire of negative regulators has therefore evolved, acting through deubiquitination, dephosphorylation, proteasomal degradation, sequestration, or competitive binding (9, 21–25). Identifying the full set of these brakes and the molecular interfaces through which they operate is a prerequisite for any therapeutic strategy that seeks to recalibrate rather than simply suppress antiviral signaling.

Members of the interleukin-17 (IL-17) receptor family are increasingly recognized as regulators of innate immune signaling beyond their canonical cytokine axis (26, 27). IL-17RD, also known as Similar Expression to Fgf genes (SEF), was originally identified in zebrafish as a feedback antagonist of FGF-stimulated Ras/MAPK signaling (28, 29) and is frequently downregulated in human carcinomas, where its loss promotes tumor progression and inflammation-associated tumorigenesis (30–35). IL-17RD remains an orphan receptor, as no IL-17 family ligand has been assigned to it. Nonetheless, it contains the cytoplasmic SEFIR domain that defines the family, and Mellett and colleagues demonstrated that IL-17RD dampens Toll-like receptor-driven inflammatory signaling through SEFIR/TIR-dependent interactions (36), and that it also tunes IL-17A-driven responses required for neutrophilia (37), positioning the receptor as a node that integrates mitogenic and innate immune inputs. Whether IL-17RD also regulates cytosolic RNA sensing, a distinct sensing modality that converges on MAVS rather than on MyD88 or TRIF, has not been examined.

Here, we report that IL-17RD is a negative regulator of the RLR pathway. Using loss- and gain-of-function studies in a human airway epithelial carcinoma cell line, epistasis analysis, subcellular localization, co-immunoprecipitation, and size-exclusion fractionation of native signalosomes, as well as infection of *Il17rd*^−/-^ mice with SARS-CoV-2, we show that IL-17RD acts at the MAVS platform on the ERGIC to restrain both the IRF3 and NF-κB arms of the antiviral response, and that its intracellular TIR subdomain is sufficient for this activity. Loss of this brake amplifies antiviral and inflammatory output and, *in vivo*, aggravates virus-induced lung pathology.

## RESULTS

### IL-17RD negatively regulates the RLR-dependent TBK1-IRF3 signaling cascade

Transcriptomic profiling of spleens from *Il17rd*^−/-^ mice provided the initial indication that IL-17RD constrains antiviral gene expression. Gene Ontology and KEGG enrichment analyses of significantly upregulated differentially expressed transcripts identified antiviral and IFN response signatures as the dominant categories (**Supplemental Figure 1**). To test this directly in a human cell system relevant to respiratory viral infection, we generated IL-17RD-deficient A549 cells by CRISPR/Cas9 editing and, independently, depleted the receptor with shRNA. Two independently edited clones were validated by sequencing and RT-qPCR (**Figure 1A-B**; **Supplemental Figure 2A-B**), and shRNA-expressing populations showed efficient reduction of *IL17RD* mRNA (**Figure 1C**; **Supplemental Figure 2C**). We then monitored the activation state of the TBK1-IRF3 module. In WT cells, poly I:C transfection, EMCV infection, and SeV infection each induced a transient wave of TBK1 phosphorylation followed by phosphorylation of IRF3 on Ser386. In IL-17RD-deficient clones, both phosphorylation events were markedly increased in amplitude and sustained over the 8-hour time course (**Figure 1D-F**). The same derepression was observed in shRNA-depleted populations, excluding clonal artefact (**Figure 1G-I**), and was reproduced across a second independent KO clone and three independent shRNAs (**Supplemental Figure 2D-G**). Densitometric quantification of at least four independent experiments confirmed the significance of these differences. Conversely, enforced expression of IL-17RD in 293T cells suppressed poly I:C- and SeV-induced IRF3 dimerization, as assessed by native gel electrophoresis (**Supplemental Figure 3**), demonstrating that the receptor is not merely permissive but actively antagonistic to IRF3 activation.

**Figure 1.**
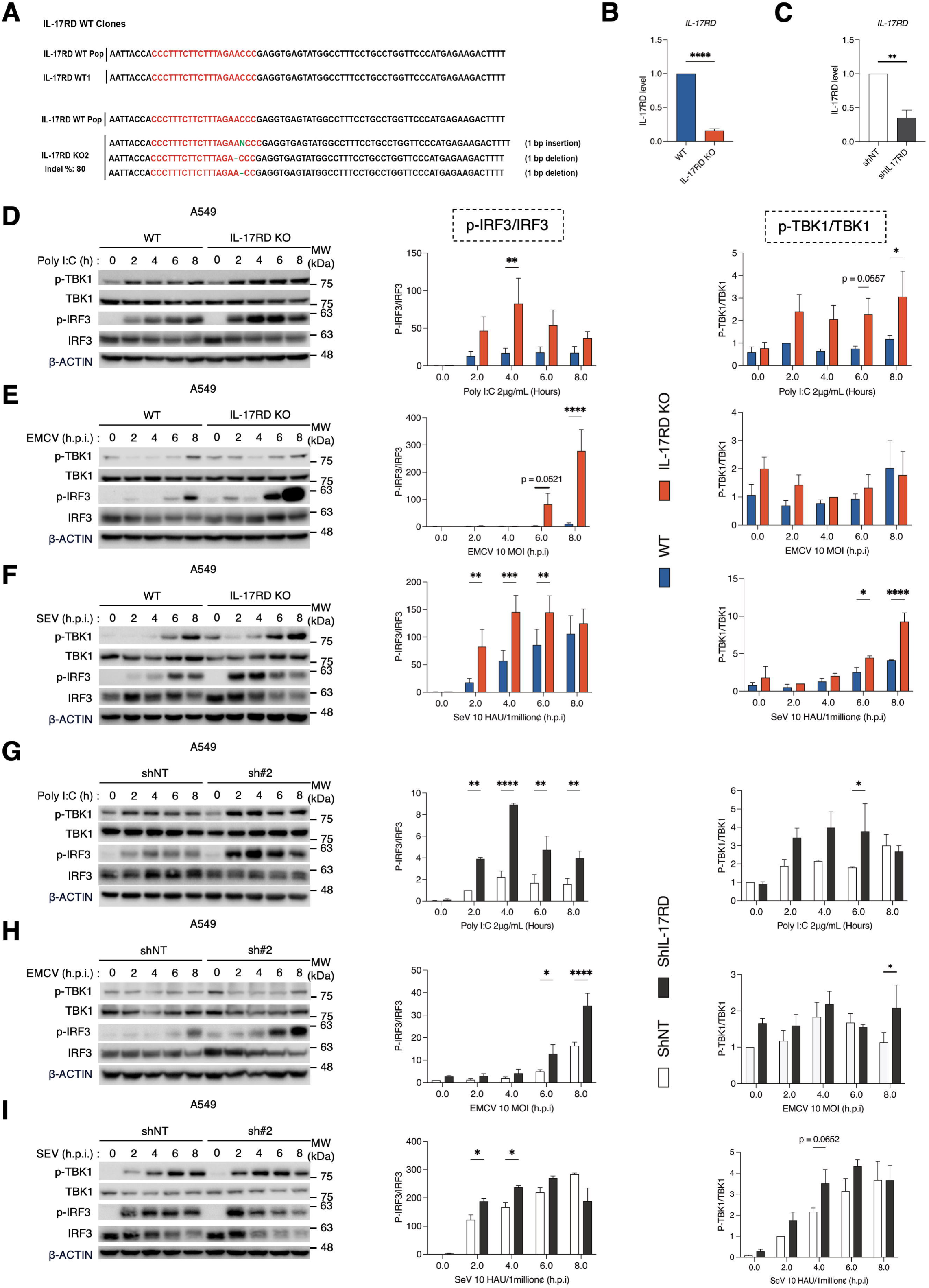
IL-17RD negatively regulates the RLR-dependent TBK1-IRF3 axis. (A) Sequencing analysis of the CRISPR-edited A549 clone #2 (KO2) aligned to the parental (WT) population and clone (WT#1); the guide RNA target sequence is shown in red. Three edited alleles were recovered (one 1-bp insertion and two 1-bp deletions; indel frequency 80%). (B) RT-qPCR analysis of IL17RD mRNA in clone KO2 relative to the WT cell population (n=3). (C) RT-qPCR analysis of IL17RD mRNA in a population of A549 cells stably expressing an shRNA (shIL-17RD, sh#2) targeting *IL17RD* mRNA, relative to a non-targeting control (shNT) (n=3). (D-F) Immunoblot analysis of A549 WT#1 and the KO2 clones transfected with poly I:C (2 µg/mL) or infected with EMCV (MOI of 10) or SeV (10 HAU/10⁶ cells) for the indicated times. (G-I) Immunoblot analysis of shNT and sh#2 A549 cell populations transfected with poly I:C or infected with EMCV or SeV for the indicated times. Immunoblot data are representative of at least four independent experiments and were quantified by densitometry as phospho-IRF3/IRF3 and phospho-TBK1/TBK1 ratios (right histograms). Data are mean ± SD. *p < 0.05, **p < 0.01, ***p < 0.001, ****p < 0.0001.

### IL-17RD also restrains the IKKβ-NF-κB and JNK1/2-p38 inflammatory signaling cascades

RLR engagement activates NF-κB and the stress-activated MAPKs in parallel with IRF3. In IL-17RD-deficient A549 cells transfected with poly I:C, phosphorylation of TAK1, IKKβ and IκBα was increased relative to parental cells, and the accompanying loss of total IκBα was accelerated (Figure 2A). Phosphorylation of JNK1, JNK2 and p38 was likewise enhanced (Figure 2B). We emphasize that these MAPK readouts are reported as parallel MAVS-dependent outputs rather than as a single linear relay. Our data establish that TAK1, JNK1/2 and p38 are more strongly phosphorylated in the absence of IL-17RD, but they do not identify the intervening MAP2K architecture, which is stimulus- and cell-type-dependent (10–13). Thus, the brake imposed by IL-17RD is not restricted to the IRF3 axis but affects the RLR output broadly, consistent with an action at or above the branch point where the MAVS platform and its TRAF-dependent scaffolds distribute the signal to the IKK-related kinases, to the TAK1-IKK module and to the stress-activated MAPKs. The functional consequence was confirmed by the translocation of transcription factors. Immunofluorescence microscopy of poly I:C-transfected cells revealed a significantly larger fraction of IL-17RD-deficient cells with predominantly nuclear IRF3 and p65 (Figure 3A-D), and the same shift was observed after EMCV infection (Figure 3E-F).

**Figure 2.**
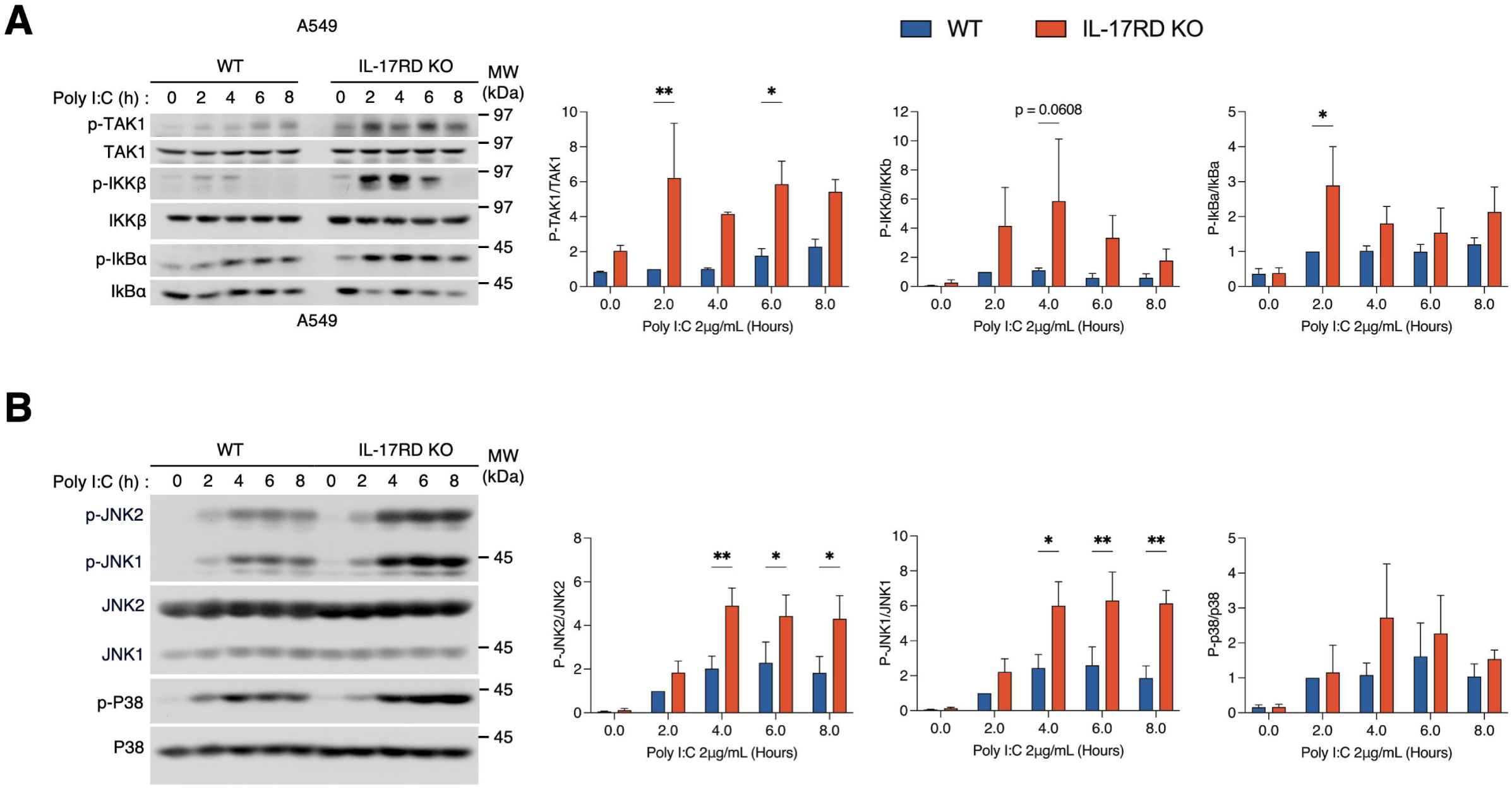
IL-17RD negatively regulates the RLR-dependent IKKβ-NF-κB and TAK1-JNK1/2-p38-AP-1 signaling cascades. (A-B) Immunoblot analysis of A549 WT#1 and the KO2 clones transfected with poly I:C (2 µg/mL) for the indicated times. Immunoblot data are representative of at least three independent experiments and were quantified by densitometry as phospho/total ratios (right histograms). Data are mean ± SD. *p < 0.05, **p < 0.01.

**Figure 3.**
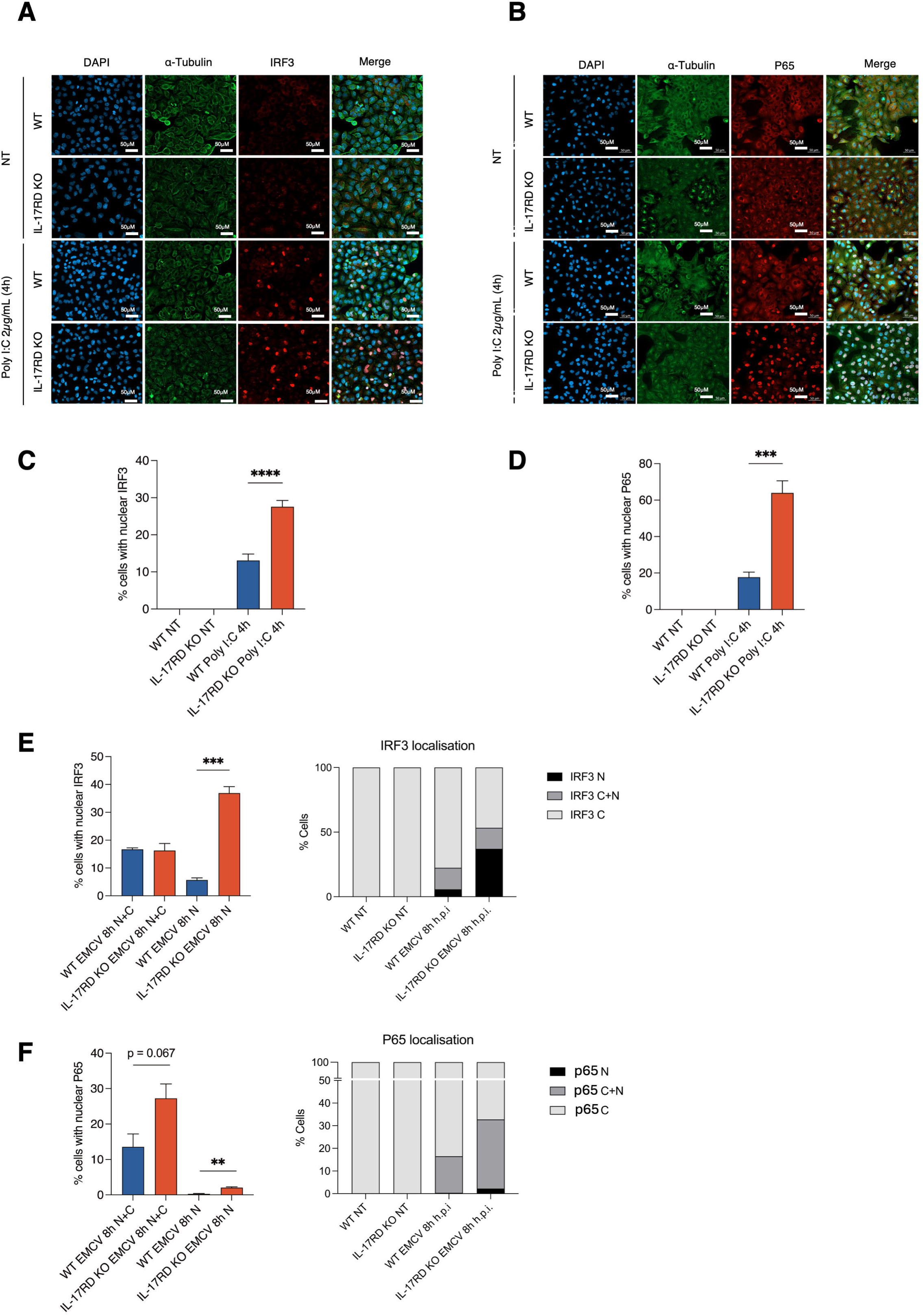
IL-17RD negatively regulates the RLR-dependent nuclear accumulation of IRF3 and p65 transcription factors. (A-B) A549 WT#1 and the IL-17RD KO clonal cell line (KO2) were left untreated (NT) or transfected with poly I:C (2 µg/mL) for 4 h, then fixed and immunostained for IRF3 (A) or p65 (B) together with α-tubulin; nuclei were labelled with DAPI and cells visualized by fluorescence microscopy. Scale bars, 50 µm. (C-D) Quantification of the percentage of cells displaying nuclear IRF3 (C) or nuclear p65 (D) under the conditions shown in (A-B). (E-F) A549 WT#1 and the KO2 clones were left untreated or infected with EMCV (MOI of 10) for 8 h and immunostained as above. The percentage of cells with nuclear (N) or mixed nuclear and cytosolic (N+C) IRF3 (E) and p65 (F) is shown (left panels), together with the full distribution scored as mostly nuclear (N), equal cytosolic and nuclear (C+N), or mostly cytosolic (C) (right panels). All quantifications are from three independent experiments (n=3). Data are mean ± SD. **p < 0.01, ***p < 0.001, ****p < 0.0001.

### Loss of IL-17RD amplifies antiviral and inflammatory gene induction

We next asked whether the augmented signaling produced a corresponding transcriptional output. IL-17RD-deficient cells transfected with poly I:C or infected with SeV or EMCV showed significantly greater induction of IFNB1, IFNL3, CCL5, IL6 and the NF-κB target NFKBIA than parental cells (Figure 4A-C). ELISA measurements of culture supernatants after 24 hours showed higher IL-6 and IFN-β in the absence of IL-17RD (Figure 4D). Notably, the magnitude of derepression was greatest for EMCV, the MDA5-dependent stimulus, and for the type III IFN genes, which are the dominant IFN species produced by airway epithelium. To assess relevance to a clinically important pathogen, we silenced IL-17RD in A549 cells stably expressing human ACE2 and then infected them with SARS-CoV-2. Depletion of IL-17RD strongly potentiated the induction of IFNB1, IFNL1, IFNL2, IFNL3, ISG15, ISG56, IRF7, CCL5 and IL6 at a multiplicity of infection of 2 (Figure 5), and the same pattern was observed at a lower multiplicity of 0.1 (Supplemental Figure 4). SARS-CoV-2 is known to be a comparatively weak inducer of IFN (19, 38–43); the observation that removing a single host brake substantially restores this induction identifies IL-17RD as a limiting factor in the response to this virus.

**Figure 4.**
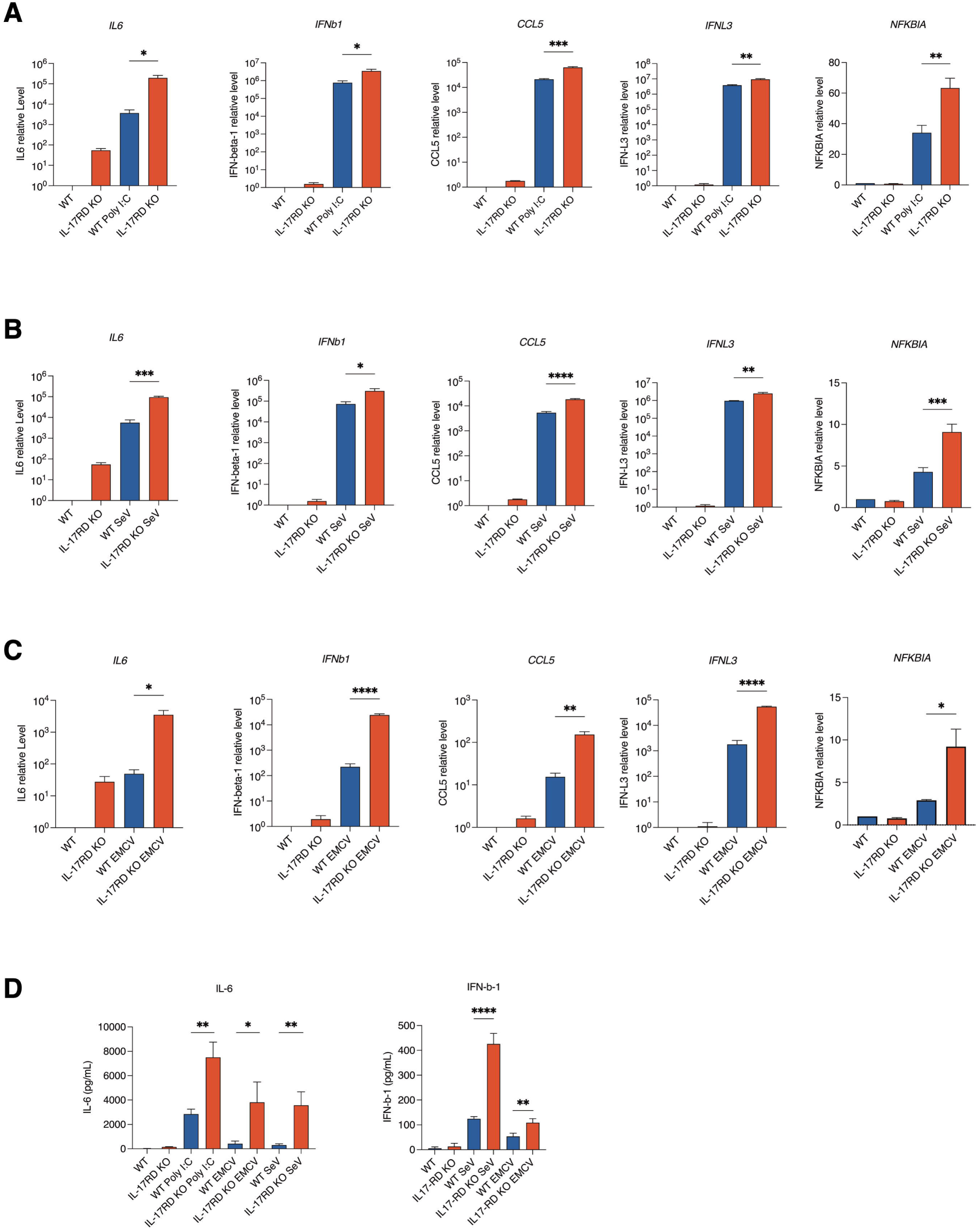
IL-17RD regulates the induction of antiviral and inflammatory cytokines. The parental A549 WT#1 clone (WT; blue bars) and the edited A549 clone (KO2; red bars) were left untreated or transfected with poly I:C (2 µg/mL) for 4 h (A), or infected with SeV (10 HAU/10⁶ cells) for 8 h (B) or EMCV (MOI of 10) for 8 h (C). Total RNA was isolated and analyzed by RT-qPCR for *IL6, IFNB1, CCL5, IFNL3* and *NFKBIA* (n=3; note the logarithmic scale for IL6, IFNB1, CCL5 and IFNL3). (D) Cells were treated or infected as above for 24 h and supernatants were collected for measurement of secreted IL-6 and IFN-β by ELISA (n=3). Data are mean ± SD. *p < 0.05, **p < 0.01, ***p < 0.001, ****p < 0.0001.

**Figure 5.**
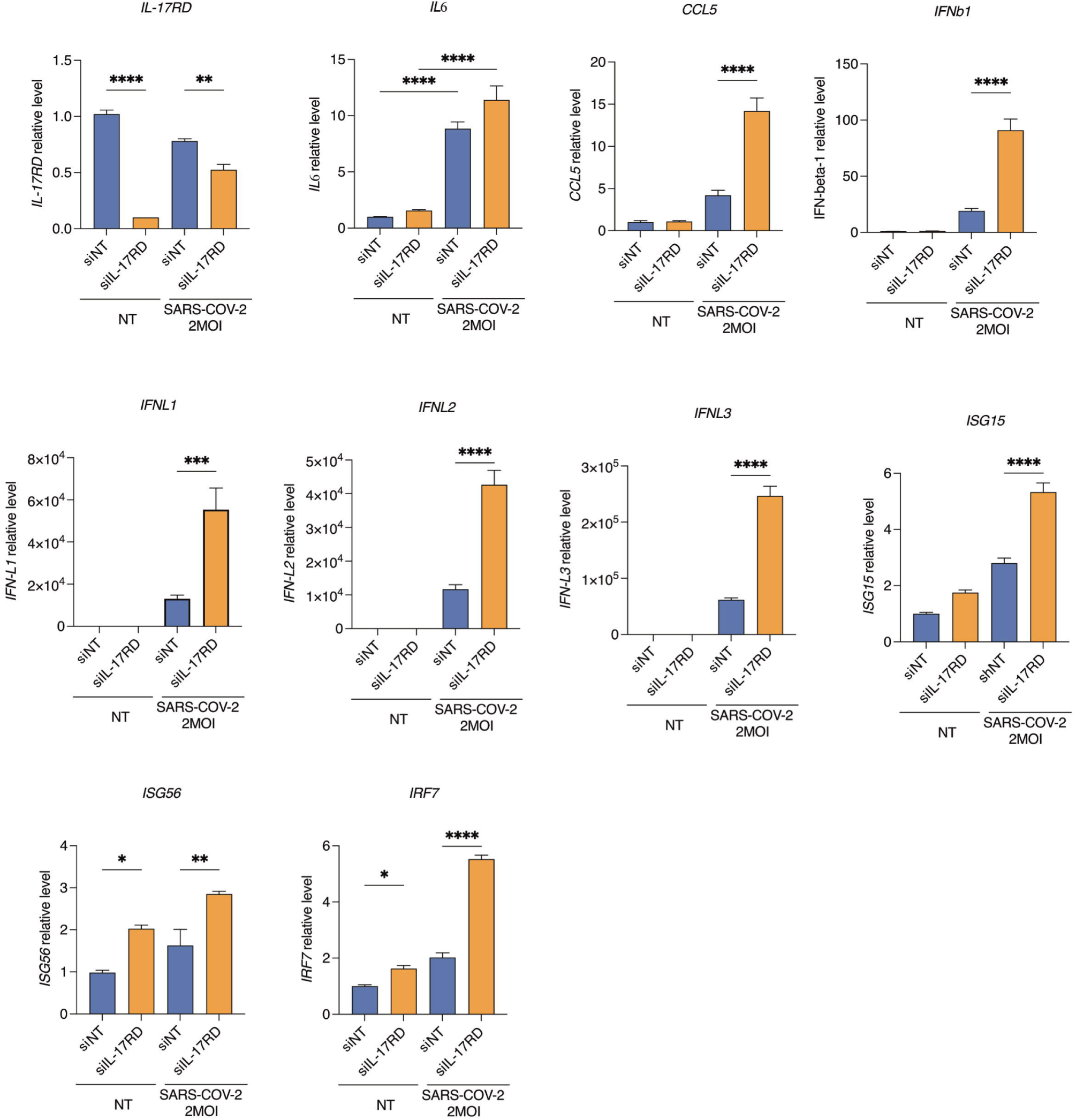
IL-17RD regulates the induction of antiviral and inflammatory cytokines in SARS-CoV-2-infected cells. A549 cells stably expressing the human ACE2 receptor were transfected with control (siNT) or IL17RD pooled siRNAs (siIL-17RD) for 72 h and then left untreated (NT) or infected with SARS-CoV-2 (MOI of 2) for 24 h as indicated. Total RNA was isolated and analyzed by RT-qPCR for *IL17RD, IL6, CCL5, IFNB1, IFNL1, IFNL2, IFNL3, ISG15, ISG56* and *IRF7* (n=3). Data are mean ± SD. *p < 0.05, **p < 0.01, ***p < 0.001, ****p < 0.0001.

### IL-17RD acts at the level of MAVS, downstream of the RLR sentinels

To position IL-17RD within the cascade, we performed epistasis experiments in 293T cells using ISRE- and NF-κB-driven luciferase reporters alongside expression constructs for successive pathway components. Increasing amounts of IL-17RD dose-dependently suppressed ISRE and NF-κB reporter activation induced by RIG-I, MDA5, and TRIF, consistent with the reported activity of the receptor in TLR3 signaling (36) (Supplemental Figure 5A-B). Inhibition was preserved when the pathway was driven by MAVS overexpression but was lost when driven by TBK1 or the constitutively active IRF3-5D mutant (Supplemental Figure 5C-D). The point of action of IL-17RD therefore lies at the level of the MAVS platform, downstream of the sensors and upstream of the TBK1-IRF3 module.

### IL-17RD localizes to the ERGIC and associates with RLR effectors

Our laboratory previously identified ER-to-Golgi trafficking components as essential to the antiviral response (4, 44), showing that TRAF3 co-exists with MAVS in the Endoplasmic-Reticulum-Golgi Intermediate Compartment (ERGIC). As a transmembrane receptor, IL-17RD could also be in close proximity to the MAVS signalosome. We stably complemented the IL-17RD KO clone with either empty vector or MYC-tagged full-length IL-17RD (Figure 6A). Confocal microscopy revealed extensive colocalization of MYC-IL-17RD with the ERGIC marker ERGIC53, with a mean Pearson correlation coefficient of approximately 0.8, well above the 0.5 threshold (Figure 6B-C). We then tested for physical association with pathway components. Co-immunoprecipitation from 293T extracts showed that MYC-IL-17RD associates with FLAG-tagged RIG-I, MDA5, MAVS, TBK1 and IRF3, but not with FLAG-GFP (Figure 6D). Densitometric quantification across five independent experiments showed the strongest recovery with MDA5, followed by MAVS and IRF3, with weaker but reproducible association with RIG-I and TBK1 (Figure 6E). IL-17RD therefore resides in an intracellular compartment where it negatively regulates the effectors of the antiviral signalling pathway.

**Figure 6.**
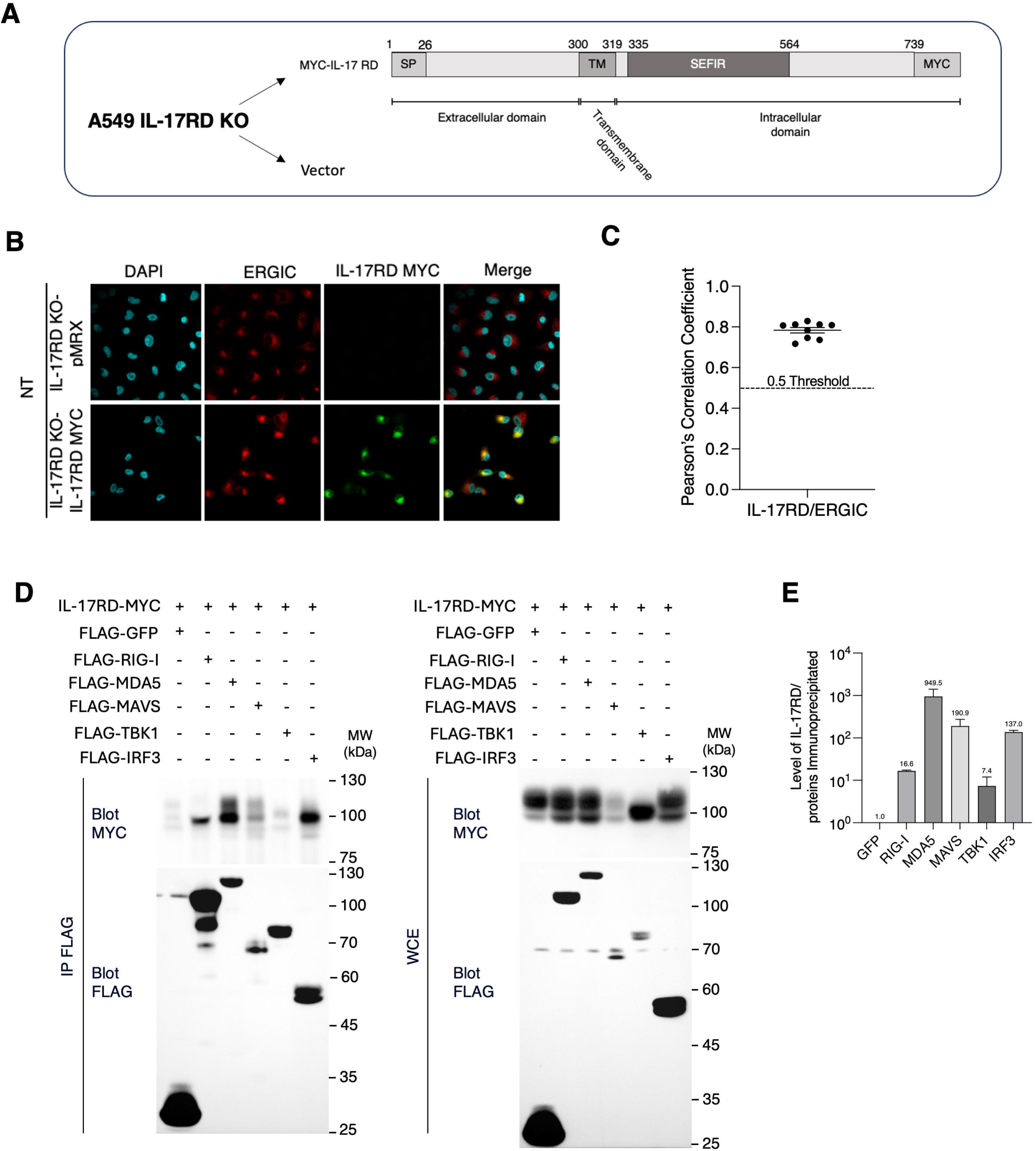
IL-17RD is preferentially expressed at the ERGIC and associates with RLR effectors. (A) Schematic of the stable complementation of the A549 IL-17RD KO cell line (KO2) with either the empty vector (pMRX) or MYC-tagged full-length IL-17RD (IL-17RD KO-IL-17RD MYC). Residue numbering delineates the signal peptide (1–26), the extracellular domain (27–300), the transmembrane segment (301–319), the intracellular domain containing the SEFIR domain (335–564) and the C-terminal MYC tag (starting at residue 740). (B) The indicated cell lines were left untreated (NT), fixed, permeabilized and immunostained with anti-MYC and anti-ERGIC53 antibodies. Nuclei were labelled with DAPI and cells visualized by confocal microscopy. Images are representative of four observations. (C) Pearson’s correlation coefficient was used to quantify colocalization between MYC-tagged IL-17RD and ERGIC53; the dashed line indicates the 0.5 threshold (n=9 fields of view). (D) Co-immunoprecipitation experiments performed on whole cell extracts (WCE) of 293T cells previously transfected with the indicated constructs. Anti-FLAG immunoprecipitates (left) and matched WCE (right) were analyzed by immunoblotting with anti-MYC and anti-FLAG antibodies. Immunoblots are from a single experiment and are representative of five independent experiments. (E) Immunoblot data were quantified by densitometry and expressed as the ratio of MYC-IL-17RD to the corresponding immunoprecipitated protein, normalized to the FLAG-GFP control (set to 1.0).

### IL-17RD re-expression remodels high-molecular-weight signalosome fractions

If IL-17RD antagonizes signalosome function, its presence should alter the composition of native high-molecular-weight MAVS signalosome complexes co-eluting around 600 to 700 kDa (45–47). We subjected extracts from the complemented KO lines to size-exclusion chromatography. MYC-IL-17RD eluted predominantly in fractions 4-5, corresponding to a mass of approximately 670 kDa, and its presence clearly shifted the distribution of TRAF3, with more modest changes in TRAF6, MDA5 and MAVS, across the gradient (Figure 7A). We therefore isolated fraction 5 from vector- and IL-17RD-complemented cells, untreated or transfected with poly I:C for 3 hours, and analyzed its composition (Figure 7B). In vector-complemented cells, poly I:C stimulation loaded the ∼670 kDa complex with IRF3, while TRAF6, RIG-I and, to a lesser extent, MDA5 remained detectable. Re-expression of IL-17RD reduced the amount of TRAF6 and MDA5 recovered in this fraction and, most strikingly, curtailed the stimulus-dependent recruitment of IRF3, while TBK1 and RIG-I levels were comparatively unaffected. IL-17RD thus acts as a component of the ∼670 kDa signalosome that limits the productive assembly of effectors within it, particularly the delivery of the IRF3 substrate to the complex. This selective effect on MDA5 and IRF3 parallels their co-immunoprecipitation with the receptor (Figure 6D-E), indicating that the association of IL-17RD with these two effectors is reflected in their redistribution away from the productive complex.

**Figure 7.**
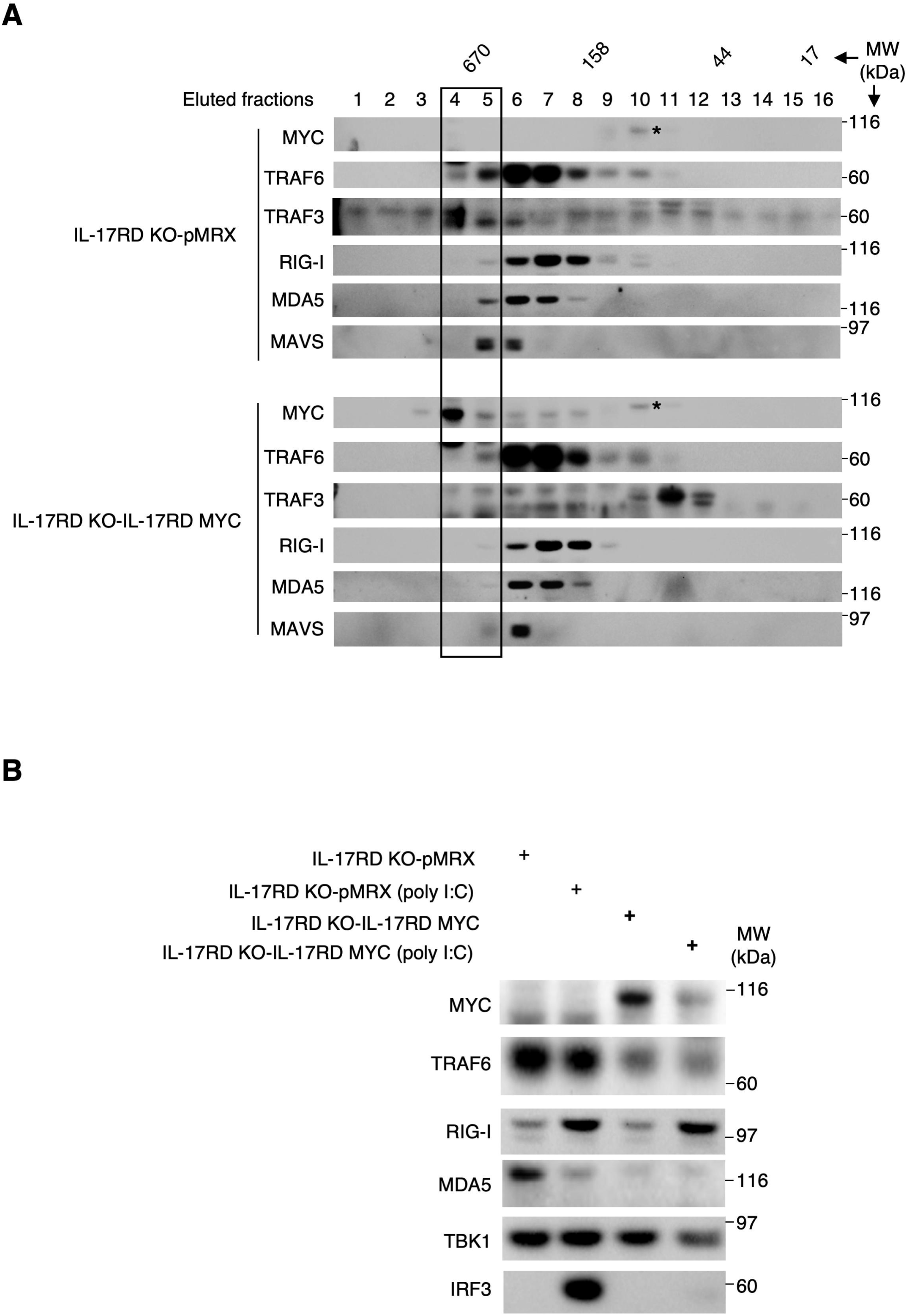
IL-17RD re-expression alters the distribution of RLR effectors across the high-molecular-weight signalosome fractions. (A) Whole cell extracts from the IL-17RD KO clonal cell line (KO2) complemented with empty vector (IL-17RD KO-pMRX) or with MYC-tagged IL-17RD (IL-17RD KO-IL-17RD MYC) were resolved by size-exclusion chromatography. The sixteen collected fractions were analyzed by immunoblotting with the indicated antibodies. The elution positions of the 670, 158, 44 and 17 kDa calibration standards are indicated, and the boxed lane marks fraction 4-5 where the complemented expression of the MYC-tagged receptor is mostly observed. (B) The same complemented cell lines were left untreated or transfected with poly I:C (2 µg/mL) for 3 h. Whole cell extracts were resolved by size-exclusion chromatography and fraction 5 was analyzed by immunoblotting using the indicated antibodies. Data are representative of three independent experiments. * Non-specific signal.

### The intracellular TIR subdomain is sufficient for antagonistic activity

IL-17RD is a single-pass transmembrane protein comprising an extracellular region (residues 27-300), a transmembrane segment (residues 300-319), and an intracellular region containing the SEFIR domain (residues 335-564), within which the TIR subdomain spans residues 355-508 (Figure 8A). Because SEFIR/TIR interactions mediate the inhibitory action of IL-17RD on TLR signaling (36), we asked whether this module alone is sufficient in the RLR context. We complemented the KO clone with MYC-tagged full-length IL-17RD or with the MYC-tagged TIR subdomain alone. Both constructs suppressed poly I:C-induced IRF3 phosphorylation, and the TIR subdomain was at least as effective as the full-length receptor (Figure 8B). Consistent with this, both restored repression of CCL5, IFNB1, and IFNL3 induction (Figure 8C). The intracellular TIR subdomain therefore constitutes the minimal functional unit of the brake, and neither the extracellular domain nor the receptor’s FGF-antagonistic function is required for its antiviral inhibitory activity.

**Figure 8.**
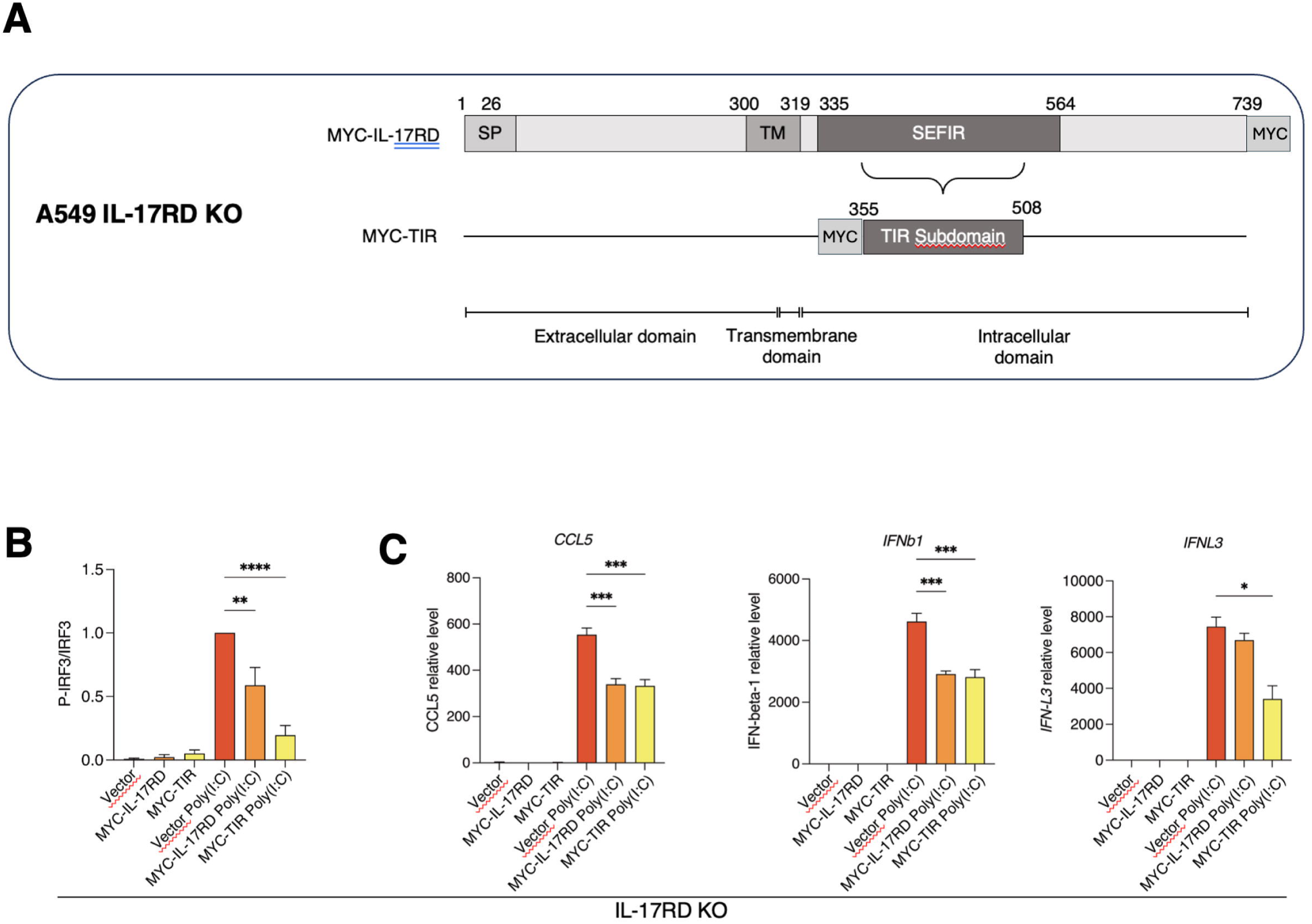
Complementation studies in the IL-17RD KO clone indicate that the TIR subdomain of IL-17RD is sufficient to confer antagonistic activity. (A) Schematic of the stable complementation of the A549 IL-17RD KO cell line (KO2) with either MYC-tagged full-length IL-17RD (MYC-IL-17RD) or the MYC-tagged TIR subdomain alone (MYC-TIR; residues 355-508). (B-C) The indicated cell lines were left untreated or transfected with poly I:C (2 µg/mL) for 3 h. (B) Cellular extracts were analyzed by immunoblotting with anti-phospho-IRF3 (Ser386) and anti-IRF3 antibodies; data were quantified by densitometry as the phospho-IRF3/IRF3 ratio (n=3). (C) Total RNA was isolated and analyzed by RT-qPCR for *CCL5, IFNB1* and *IFNL3* (n=3). Data are mean ± SD. *p < 0.05, **p < 0.01, ***p < 0.001, ****p < 0.0001.

### Loss of IL-17RD aggravates SARS-CoV-2-induced lung pathology in vivo

To determine whether this brake operates in an intact animal, we turned to *Il17rd*^−/-^ mice. Primary lung fibroblasts isolated from these animals showed hyper-induction of *Ifnb1* and the interferon-stimulated gene *Rsad2* after poly I:C transfection compared with wild-type cells (**Figure 9A**), demonstrating that the cell-intrinsic phenotype observed in human cells is conserved in primary murine cells. *Il17rd*^+/+^ and *Il17rd*^−/-^ mice were then infected intranasally with a moderate dose of the SARS-CoV-2 beta variant (35,000 TCID_50_) and followed for 10 days (**Figure 9B**). Body weight did not differ significantly between genotypes over this period (**Figure 9C**), and lung viral titres, measured as TCID_50_ on days 3, 5 and 10 post-infection, were comparable (**Figure 9D**). Despite equivalent viral burden, pulmonary cytokine levels diverged: IFN-α, IFN-β, IFN-γ and TNF-α were significantly elevated in *Il17rd*^−/-^ lungs at day 5 post-infection, the peak of the response (**Figure 9E**). The phenotype is therefore one of dysregulated response magnitude rather than altered viral control. Histopathological analysis established the consequences of this dysregulation. At day 5 post-infection, *Il17rd*^−/-^ lungs showed increased atelectasis and prominent peribronchiolar and perivascular inflammatory infiltrates (**Figure 10A**), resulting in significantly higher fibrosis and inflammation scores than in infected wild-type animals (**Figure 10B-C**). IL-17RD therefore protects the lung from virus-induced inflammatory damage without compromising viral clearance.

**Figure 9.**
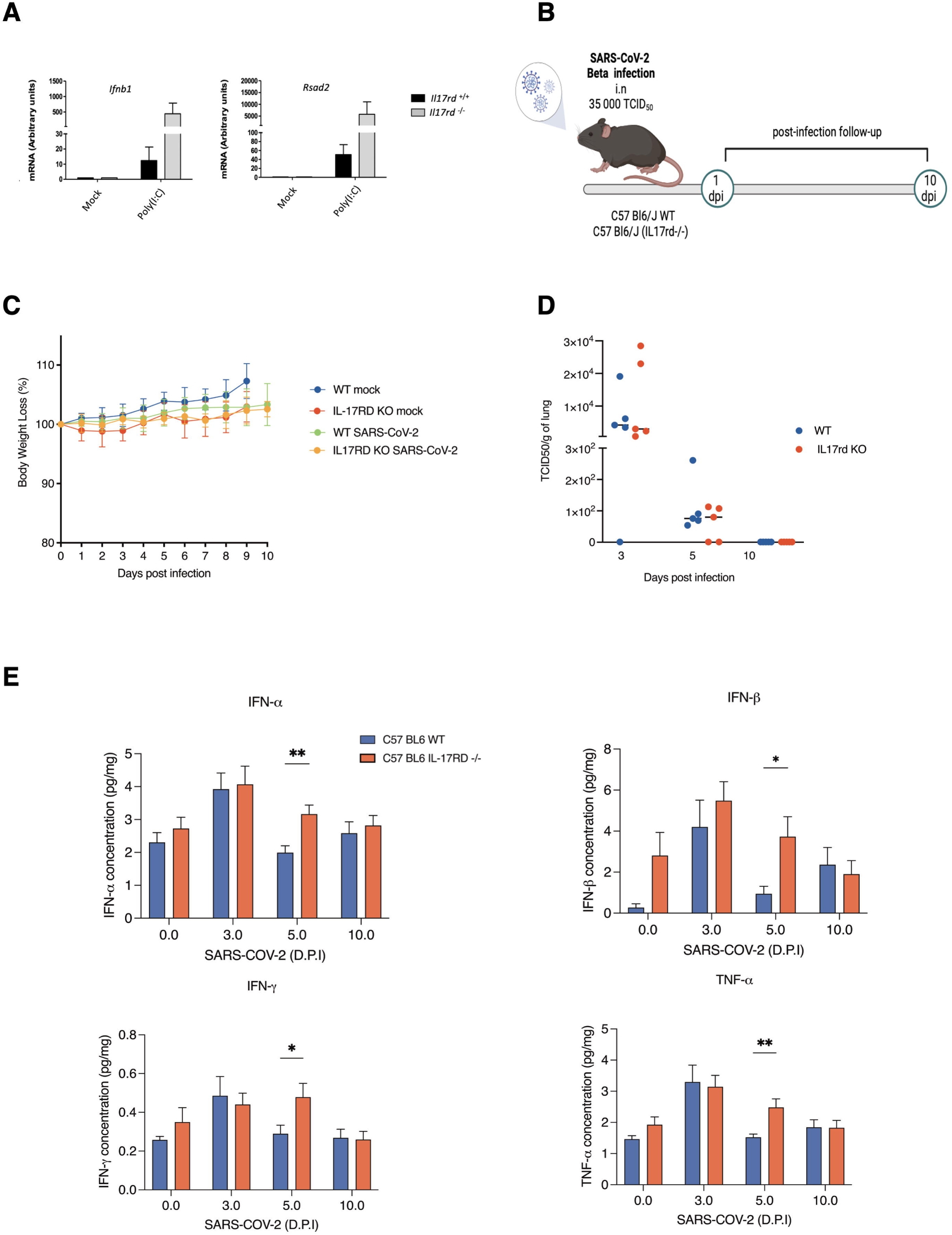
IL-17RD controls the magnitude of the inflammatory response to a moderate dose of SARS-CoV-2 in vivo. (A) Primary lung fibroblasts were isolated from *Il17rd*^+/+^ (black bars) and *Il17rd*^−/-^ (grey bars) mice, amplified in culture and left untreated (mock) or transfected with poly I:C (1 µg/mL) for 6 h. Total RNA was purified and analyzed by RT-qPCR for *Ifnb1* and *Rsad2* (n=3). (B) Schematic of the infection protocol: C57BL/6J wild-type (*Il17rd*^+/+^) and *Il17rd*^−/-^ mice received a moderate intranasal dose of SARS-CoV-2 beta variant (35,000 TCID₅₀) and were followed for 10 days post-infection. (C) Body weight of mock- and SARS-CoV-2-infected animals of both genotypes, expressed as a percentage of the day-0 value, over the 10-day period. (D) Viral titre in the lungs of infected *Il17rd*^+/+^ and *Il17rd*^−/-^ mice, measured as the 50% tissue culture infectious dose (TCID₅₀) per gram of lung at 3, 5 and 10 days post-infection (n=5 per genotype and time point). (E) IFN-α, IFN-β, IFN-γ and TNF-α concentrations (pg/mg) in lung homogenates from wild-type (blue bars) and *Il17rd*^−/-^ (orange bars) mice at 0, 3, 5 and 10 days post-infection (n=5). Data are mean ± SD. *p < 0.05, **p < 0.01.

**Figure 10.**
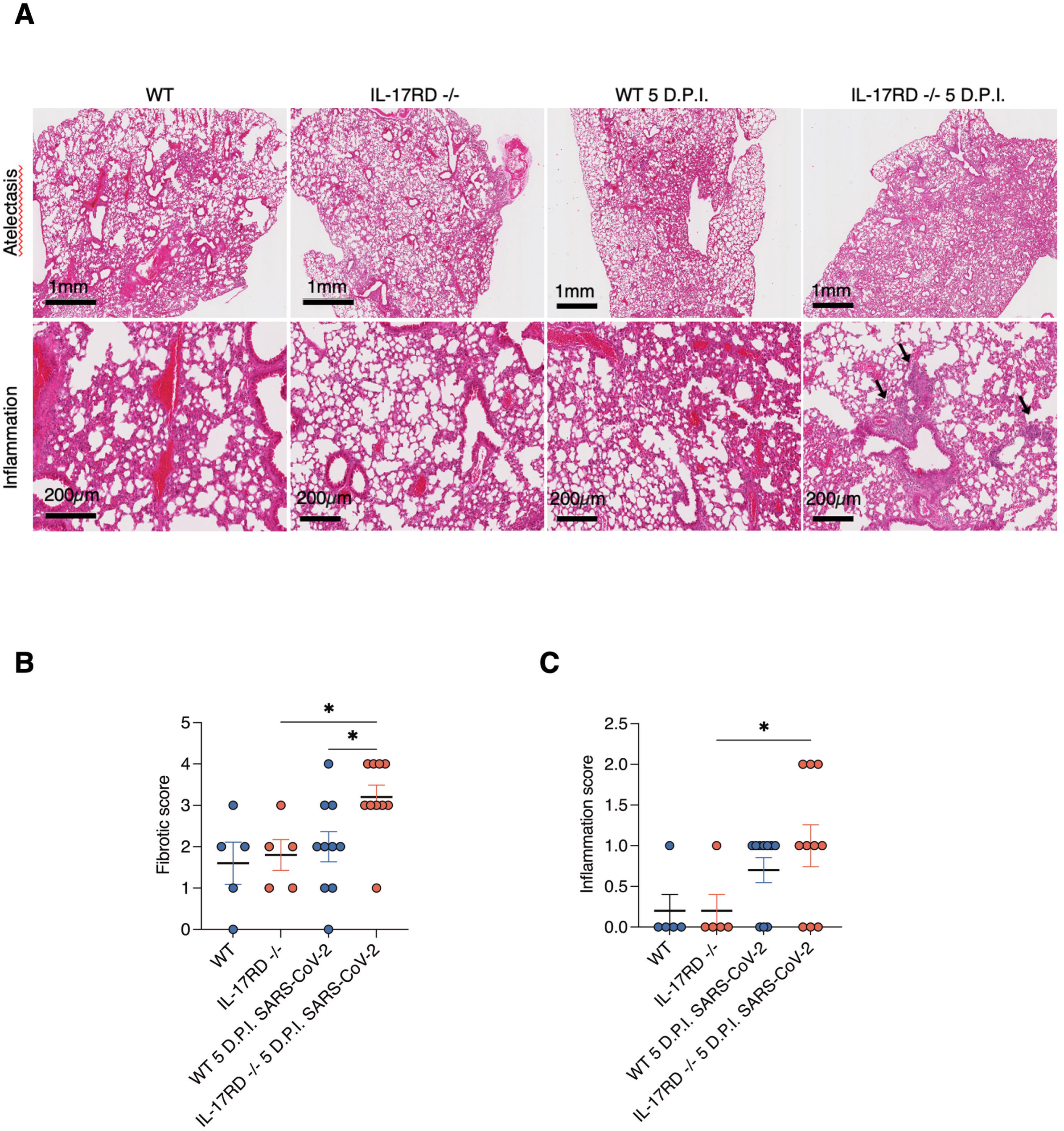
IL-17RD protects from lung inflammation and fibrosis in response to a moderate dose of SARS-CoV-2 in vivo. (A) Representative eosin-stained lung sections from uninfected wild-type (*Il17rd*^+/+^) and *Il17rd*^−/-^ mice and from mice at 5 days post-infection (D.P.I.) with SARS-CoV-2. Upper row, low magnification showing atelectasis (scale bars, 1 mm); lower row, higher magnification showing inflammatory infiltrates (scale bars, 200 µm). Arrowheads indicate peribronchiolar and perivascular inflammatory foci. (B-C) Fibrotic (B) and inflammation (C) scores assigned to lung lesions by a board-certified veterinary anatomic pathologist blinded to genotype (n=5 to n=10 as shown). Data are mean ± SD. *p < 0.05.

## DISCUSSION

This study identifies IL-17RD as a negative regulator of the RLR-MAVS axis and, more broadly, extends the regulatory reach of the IL-17 receptor family into cytosolic nucleic acid sensing. Four lines of evidence support this conclusion. First, loss of IL-17RD in the human airway epithelial-derived lung carcinoma A549 cell line, achieved independently by CRISPR editing and shRNA depletion, amplifies and prolongs activation of the TBK1-IRF3 and IKKβ-NF-κB modules, as well as the JNK1/2 and p38 MAPK branches, and increases the resulting antiviral and inflammatory transcriptional output, including in response to SARS-CoV-2. Second, enforced expression of IL-17RD suppresses IRF3 dimerization, establishing an actively antagonistic relationship. Third, IL-17RD localizes to the ERGIC, associates with RIG-I, MDA5, MAVS, TBK1, and IRF3, and its presence remodels the composition of native ∼670 kDa signalosome fractions, thereby reducing IRF3 delivery to the complex. Fourth, *Il17rd*^−/-^ mice mount an exaggerated pulmonary cytokine response to SARS-CoV-2 and sustain greater lung inflammation and fibrosis at an equivalent viral load.

The epistasis data place the point of action at MAVS, and the fractionation data suggest a mechanism based on signalosome composition rather than catalytic inhibition. IL-17RD co-elutes with the high-molecular-weight complex, and when present, the complex recovers less TRAF3, TRAF6 and MDA5, and markedly less stimulus-dependent IRF3, while TBK1 abundance is largely preserved. A signalosome that retains its kinase but is depleted of substrate offers a straightforward explanation for the reduced IRF3 phosphorylation observed in complemented cells and for the mirror-image hyperphosphorylation observed when IL-17RD is absent. This model is consistent with the accepted architecture of the MAVS platform, in which TRAF-dependent ubiquitin scaffolds govern the stoichiometry of kinase and substrate recruitment (8, 21), and with our earlier demonstration that the TRAF3 interactome links ER-to-Golgi compartments to innate immune output (4).

That the isolated TIR subdomain reproduces, and if anything exceeds, the inhibitory activity of the full-length receptor is informative in two respects. Mechanistically, it indicates that the brake is likely executed by protein-protein interaction through the intracellular SEFIR/TIR subdomain module, consistent with the SEFIR/TIR-dependent mechanism established for IL-17RD in TLR signaling (36). It also shows that the antiviral brake requires neither the ectodomain nor the transmembrane region, which have been reported to be the predominant contributors to FGF/ERK antagonism in mammalian cells (48, 49). The two activities therefore have distinct domain requirements, although the intracellular region has also been implicated in FGF antagonism and the underlying mechanism remains debated (35, 49). Practically, it defines a compact interaction surface that could be targeted pharmacologically, and it provides the basis for separation-of-function alleles that would allow the antiviral brake to be genetically disabled without perturbing the receptor’s other physiological roles.

The SEFIR domain is the defining feature of the IL-17 receptor family and mediates recruitment of the adaptor ACT1 (CIKS/TRAF3IP2) in canonical IL-17 signaling (26, 27, 50). ACT1 is a U-box E3 ubiquitin ligase that promotes K63-linked ubiquitination of TRAF6 (51), is itself regulated by TBK1- and IKKε-dependent phosphorylation (52), and has been reported to participate in antiviral signaling in an evolutionarily conserved manner (53). Our identification of a SEFIR/TIR-dependent brake acting at the MAVS signalosome therefore raises the question of which SEFIR-domain partners transmit this activity, and whether the ubiquitin-dependent scaffolds shared between the IL-17 and RLR pathways provide the point of convergence. Addressing this question will require systematic mapping of the interface and genetic dissection of candidate effectors.

The in vivo phenotype has implications for the pathophysiology of respiratory viral disease. In Il17rd-/- animals, cytokine levels and histological damage increase, whereas viral titres do not decline, indicating that the additional IFN and inflammatory output is not translated into improved viral control in this model. This dissociation is characteristic of the immunopathological component of severe respiratory infection, in which outcome correlates more closely with the intensity of the host response than with viral burden (16–20). It also implies that IL-17RD occupies a position where pharmacological modulation could be titrated in either direction: transient relief of the brake early in infection to strengthen the IFN response, particularly relevant given the weak IFN induction elicited by SARS-CoV-2 (19, 38–43), or reinforcement of the brake in the later inflammatory phase. The frequent downregulation of IL-17RD in human carcinomas (32–34) suggests that variation in receptor expression may also contribute to inter-individual differences in inflammatory tone.

Several limitations should be noted. Our cellular work relies primarily on the human airway epithelial-derived lung carcinoma A549 cell line and 293T cells, and confirmation in primary human airway epithelium and in myeloid populations will be important. The mouse experiments used a moderate infectious dose on a C57BL/6 background, where SARS-CoV-2 replication is limited. Whether the same dissociation between viral load and pathology holds in more permissive models, such as K18-hACE2 transgenic animals, remains to be established. Although the fractionation data indicate that IL-17RD alters signalosome composition, the precise stoichiometry and the identity of the direct binding partner within the complex are not resolved by the present experiments.

In summary, we establish IL-17RD as a host-encoded brake on RLR-dependent antiviral innate immunity, acting at the ERGIC-associated MAVS signalosome via its intracellular TIR subdomain, thereby protecting the lung from virus-induced inflammatory injury. These findings link an orphan receptor of the IL-17 family, previously studied in the contexts of FGF signaling, TLR responses and cancer, to the regulation of cytosolic RNA sensing, and identify a defined molecular module as a candidate point of therapeutic intervention in severe respiratory viral disease.

## MATERIALS AND METHODS

### Cell lines and culture

Human lung carcinoma A549 cells and 293T cells were obtained from the American Type Culture Collection (ATCC, Manassas, VA, USA) and maintained in DMEM supplemented with heat-inactivated 10% fetal bovine serum, 2 mM L-glutamine, and antibiotics at 37 °C in 5% CO2. The PLAT-A retroviral packaging cell line was purchased from Cell Biolabs. All cell lines were maintained according to ATCC guidelines. A549 cells stably expressing human ACE2 were generated by retroviral transduction and selected with the appropriate antibiotic. Complemented lines were produced by transduction with pMRX-based retroviral vectors encoding MYC-tagged full-length IL-17RD, the MYC-tagged TIR subdomain (residues 355-508), or an empty vector, followed by antibiotic selection. Cells were used at low passage and routinely tested for mycoplasma contamination.

### Plasmids and expression constructs

The pMRX retroviral vector, NF-κB- and ISRE-driven luciferase reporter constructs, and expression plasmids encoding TRIF, RIG-I, MDA5, MAVS, TBK1, IRF3, and the constitutively active IRF3-5D mutant have been described in our previous publications (44, 54, 55). The pMRX-IL-17RD-MYC construct was generated from the human SEF isoform a (hSEF-a) cDNA described previously (31) by inserting the coding sequence in frame with a C-terminal MYC epitope tag. The pMRX-TIR-MYC construct, encoding the isolated TIR subdomain (residues 355-508), was derived from the same template. All constructs were verified by Sanger sequencing. Retroviral particles were produced in PLAT-A packaging cells exactly as described in our recent work (56). Briefly, PLAT-A cells were transfected with 12 µg of the relevant pMRX plasmid using polyethylenimine (PEI, 3:1 PEI:DNA ratio). After 48 h, the virus-containing medium was filtered through a 0.45 µm filter (Millipore) and supplemented with 8 µg/mL polybrene (Sigma-Aldrich). Target cells were incubated with the viral supernatant for 16 h, and infected populations were subsequently enriched by antibiotic selection for 10 days to generate stable cell populations.

### CRISPR/Cas9 editing and RNA interference

A549 cell pools depleted of IL-17RD by CRISPR/Cas9 gene editing were purchased from Synthego (Redwood City, CA, USA). Edited cell populations were generated by electroporating A549 cells with SpCas9 (parental population) or with sgRNA delivered as ribonucleoproteins. The sgRNA sequence used was GGGUUCUAAAGAAGAAAGGG. Single-cell clones were expanded, and the edited alleles were characterized by Sanger sequencing. Loss of transcript was confirmed by RT-qPCR. Stable knockdown populations were generated with lentiviral shRNA constructs targeting IL17RD (sh#1-sh#3) or a non-targeting control (shNT). VSVg-pseudotyped lentiviral particles were produced in 293FT cells using Lipofectamine 3000 (Thermo Fisher Scientific, L3000015) with 1.5 µg pMDLg/pRRE, 1.5 µg pRSV-REV, and 3 µg pVSV-G, together with 6 µg of the shRNA vector. Viral supernatants were harvested after 72 h and used to transduce A549 cells in the presence of 10 µg/mL polybrene, followed by puromycin selection from day 3 post-transduction (56). Transient silencing of IL17RD used ON-TARGETplus SMARTpool siRNA reagents, consisting of a pool of four independent duplexes (Dharmacon, Horizon Discovery, Lafayette, CO, USA), transfected for 72 h with the corresponding non-targeting pool (siNT) as control.

### Stimulation and viral infections

Cells were transfected with poly I:C (2 µg/mL for A549; 1 µg/mL for primary murine fibroblasts) using a cationic lipid reagent. EMCV (VR-1762; ATCC CCL-81) was used at a multiplicity of infection (MOI) of 10, and Sendai virus (Cantell strain; obtained from Specific Pathogen-Free Avian Supply (Charles River Laboratories)) at 10 HAU per 10⁶ cells. SARS-CoV-2 infections of ACE2-expressing A549 cells were performed at an MOI of 2 or 0.1 for 24 h. All work with SARS-CoV-2 was conducted under biosafety level 3 containment.

### Immunoblotting and co-immunoprecipitation

Whole-cell extracts were prepared by resuspending cells on ice for 20 min in Triton X-100 lysis buffer (1% Triton X-100, 50 mM Tris pH 7.4, 150 mM NaCl, 50 mM NaF, 5 mM EDTA, 10% glycerol, 1 mM sodium orthovanadate, 40 mM β-glycerophosphate) supplemented with protease inhibitors, followed by centrifugation and recovery of the soluble fraction. Protein concentrations were determined by the Bradford (Bio-Rad) or BCA assay according to the manufacturer’s protocol. Extracts were resolved by SDS-PAGE and transferred to nitrocellulose. Phospho-specific antibodies (against phospho-TBK1 (Ser172), phospho-IRF3 (Ser386), phospho-TAK1 (Thr187), phospho-IKKβ (Ser177), phospho-IκBα (Ser32/Ser36), phospho-JNK1/2 (Thr183/Tyr185) and phospho-p38 (Thr180/Tyr182) were obtained from Cell Signaling Technology (Danvers, MA, USA). Antibodies recognizing the corresponding total proteins (TBK1, IRF3, TAK1, IKKβ, IκBα, JNK1/2, p38), as well as those against TRAF3, TRAF6, RIG-I, MDA5, ERGIC53, α-tubulin, the MYC and FLAG epitope tags and β-actin, were obtained from Sigma-Aldrich (Merck, St. Louis, MO, USA) or Cell Signaling Technology (Danvers, MA, USA). All antibodies were used at the dilutions recommended by the suppliers. For co-immunoprecipitation, 293T cells were transiently co-transfected with MYC-IL-17RD and the indicated FLAG-tagged constructs; 1 mg of pre-cleared whole-cell extract was incubated overnight at 4 °C on a rotating wheel with 20 µL of magnetic SureBeads Protein A/G (Bio-Rad, 161-4013/161-4023) pre-coupled to 1 µg of the indicated antibody. After five washes in lysis buffer containing protease inhibitors, immune complexes were eluted in 20 µL of 2X sample buffer and analyzed by SDS-PAGE and immunoblotting alongside matched whole-cell extracts (56). Native PAGE was used to resolve IRF3 dimers. Immunoblots were quantified by densitometry.

### Size-exclusion chromatography

Whole-cell extracts from complemented KO lines, untreated or transfected with poly I:C for 3 h, were fractionated on a Superdex 200 HR 10/30 size-exclusion column (Cytiva, formerly GE Healthcare) calibrated with molecular weight standards of 670, 158, 44 and 17 kDa. Twenty-five fractions were collected, and sixteen were analyzed by immunoblotting. Fraction 5 (∼670 kDa) was selected for comparative analysis across three independent experiments.

### Immunofluorescence and confocal microscopy

Cells were seeded onto sterile glass coverslips and allowed to adhere for 24 h under standard culture conditions. Coverslips were rinsed in cold 1X PBS, fixed in 4% paraformaldehyde in 1X PBS for 15 min at room temperature, washed extensively, and permeabilized with 0.2% Triton X-100 in 1X PBS for 3 min. Cells were blocked for 15 min at room temperature in 0.1% Triton X-100 and 1% bovine serum albumin in 1X PBS, then incubated with primary antibodies against IRF3, p65 (#8242 from Cell Signaling Technology), α-tubulin (clone DM1A, Sigma-Aldrich T6199), the MYC epitope tag, or ERGIC53, each diluted to 1 µg/mL in blocking buffer. Coverslips were mounted on glass slides with ProLong Gold Antifade Mountant containing DAPI (Thermo Fisher Scientific) for nuclear staining. Confocal imaging was performed on a Leica TCS SP8, and images were analyzed in a blinded manner using LAS X Life Science Widefield software (Leica Microsystems). Nuclear accumulation of IRF3 and p65 was scored in a blinded manner into three categories (predominantly nuclear, evenly distributed, predominantly cytosolic). Colocalization of MYC-IL-17RD with ERGIC53 was quantified using the JACoP plugin in Fiji/ImageJ as the Pearson correlation coefficient (r) between fluorescence channels across nine fields of view, using automatically determined thresholds; r > 0.5 was considered indicative of significant colocalization (56).

### RT-qPCR and ELISA

Total RNA was isolated using the RNeasy Mini Kit (Qiagen, 74106) according to the manufacturer’s instructions, quantified on a NanoPhotometer (Implen, Munich, Germany), and evaluated for integrity on a 2100 Bioanalyzer (Agilent Technologies, Palo Alto, CA, USA). RNA was reverse transcribed using the Maxima First Strand cDNA Synthesis Kit with dsDNase (Thermo Fisher Scientific). Quantitative PCR assays for the indicated human and murine transcripts were designed using the Roche Universal Probe Library, and a standard curve was generated for each assay to confirm amplification efficiency between 90% and 110%. Amplification was detected on a QuantStudio 7 instrument (Thermo Fisher Scientific). All reactions were run in triplicate, and relative mRNA levels were calculated using the comparative threshold method (2-ΔΔCt), with GAPDH, HPRT and ACTB as endogenous controls (56). Secreted IL-6 and IFN-β were measured in culture supernatants collected after 24 h, and murine lung cytokines were measured in tissue homogenates, using commercial ELISA kits according to the manufacturers’ instructions.

### Luciferase reporter assays

293T cells were co-transfected with pGL4.32 (luc2P/NF-κB-RE) or pGL4.45 (luc2P/ISRE) reporter plasmids (Promega, Madison, WI, USA), a normalizing plasmid, and expression constructs for TRIF, RIG-I, MDA5, MAVS, TBK1, the constitutively active IRF3-5D mutant, or increasing amounts of human IL-17RD, as indicated. Cells were harvested 24 h post-transfection, lysed in Passive Lysis Buffer, and extracts were assayed with the Dual-Luciferase Reporter Assay kit (Promega) according to the manufacturer’s instructions. Data are expressed as firefly luciferase values divided by the internal control Renilla luciferase values, as described previously (54, 56).

### Mice and in vivo infection

*Il17rd*^−/-^ mice on the C57BL/6J background and their wild-type littermates were bred and housed under specific pathogen-free conditions. Primary lung fibroblasts were isolated by enzymatic digestion and expanded in culture. For infection, age- and sex-matched animals were inoculated intranasally with 35,000 TCID50 of the SARS-CoV-2 beta variant and monitored for 10 days, with daily weight measurements. Lungs were harvested on days 3, 5 and 10 post-infection to determine viral titre by TCID50 assay, quantify cytokines and perform histopathology. Eosin-stained sections were scored for atelectasis, inflammation and fibrosis by a board-certified veterinary anatomic pathologist blinded to genotype. All procedures were approved by the institutional animal care and use committee and conducted in accordance with the Canadian Council on Animal Care guidelines; SARS-CoV-2 infections were performed in a biosafety level 3 facility.

### Statistical analysis

Data are presented as mean ± SD from at least three independent experiments unless otherwise stated. Comparisons between two groups used the unpaired two-tailed Student t test; comparisons across multiple groups used one- or two-way analysis of variance with appropriate post hoc correction for multiple testing. Significance is indicated as *p < 0.05, **p < 0.01, ***p < 0.001, and ****p < 0.0001.

## DECLARATIONS

## Acknowledgements

We thank the members of the Servant and Meloche laboratories for helpful discussions, the personnel of the biosafety level 3 facility at the CHU de Québec Research Center. This work was supported by a Canadian Institutes for Health Research (CIHR) grant to MJS and SM.

## Author contributions

F.E.-M. performed most of the cellular and biochemical experiments, imaging, and analyzed the data. J.R. also contributed to biochemical experiments, imaging, and analyzed the data. F.D. generated most of the DNA constructs and the stable cell lines. C.G. contributed to the murine transcriptomic analysis, mouse colony work, and epistasis analysis. A.C.D.S.P.A., E.L., and I.D. performed SARS-CoV-2 infections in vitro and in vivo, viral titrations, and analysis of lung cytokines and histopathology. P.R. contributed to the design and interpretation of the interactome and microscopy experiments. B.D. contributed to the analysis and interpretation of the SARS-CoV-2 data. L.F. supervised SARS-CoV-2 infection and Biosafety Level 3 procedures. S.M. secured funding and contributed to study design and to the interpretation of the IL-17RD/SEF work. M.J.S. conceived and supervised the study, secured funding, and wrote the manuscript.

## Data availability

All data supporting the conclusions of this study are presented in the main and supplementary figures. Transcriptomic datasets and unprocessed immunoblot images are available from the corresponding author upon reasonable request.

## Supplemental figure legends

**Supplemental Figure 1. Enrichment of antiviral response signatures in the spleen of *Il17rd***^−/-^ **mice.** Total RNA was isolated from the spleens of *Il17rd*^−/-^ and *Il17rd*^+/+^ mice and subjected to transcriptomic analysis. (A) Functional classification of the significantly upregulated differentially expressed transcripts in *Il17rd*^−/-^ mice, obtained with Metascape using the Gene Ontology Biological Processes collection of gene sets. Red bars indicate p-values (−log10). (B) Functional classification as in (A), but using the KEGG pathway collection. Orange bars indicate p-values (−log10).

**Supplemental Figure 2. IL-17RD negatively regulates the RLR-dependent TBK1-IRF3 signaling cascade.** (A) Sequencing analysis of the CRISPR-edited A549 clones #1 (KO1) and #2 (KO2). (B) RT-qPCR analysis of *IL17RD* mRNA in KO1 and KO2 relative to the parental-derived WT#1 clone (n=3). (C) RT-qPCR analysis of *IL17RD* mRNA in populations of A549 cells stably expressing three different shRNAs (sh#1, sh#2, sh#3) targeting *IL17RD* mRNA, relative to a non-targeting control (shNT) (n=3). (D-E) Immunoblot analysis of the parental (WT) and IL-17RD KO clones transfected with poly I:C (2 µg/mL; D) or infected with EMCV (MOI of 10; E), quantified by densitometry (histograms). (F-G) Immunoblot analysis of the shRNA-expressing populations (shNT and sh#1 to sh#3) transfected with poly I:C (F) or infected with EMCV (G), quantified by densitometry (histograms). Data are representative of at least four independent experiments and are mean ± SD. *p < 0.05, **p < 0.01, ***p < 0.001, ****p < 0.0001.

**Supplemental Figure 3. Enforced expression of IL-17RD blocks the dimerization of IRF3 in response to poly I:C treatment or SeV infection.** 293T cells were co-transfected with FLAG-IRF3 together with MYC-tagged IL-17RD or the empty MYC vector for 24 h, and were then left untreated (NT), transfected with poly I:C, or infected with SeV for 8 h. Cellular extracts were analyzed in parallel by denaturing SDS-PAGE, with immunoblotting for FLAG-IRF3 and MYC-IL-17RD, and by native PAGE with anti-FLAG immunoblotting to resolve the monomeric and dimerized (active) forms of IRF3. Immunoblots are representative of three independent experiments.

**Supplemental Figure 4. IL-17RD regulates the induction of antiviral and inflammatory cytokines in SARS-CoV-2-infected cells at a low multiplicity of infection.** A549 cells stably expressing the human ACE2 receptor were transfected with control (siNT) or IL17RD pooled siRNAs (siIL-17RD) for 72 h and then left untreated (NT) or infected with SARS-CoV-2 (MOI of 0.1) for 24 h as indicated. Total RNA was isolated and analyzed by RT-qPCR for the indicated transcripts (n=3). Data are mean ± SD. *p < 0.05, **p < 0.01, ***p < 0.001, ****p < 0.0001.

**Supplemental Figure 5. Epistasis analysis reveals that IL-17RD acts downstream of the RLR sentinels, at the level of MAVS.** 293T cells were co-transfected with the luciferase reporter plasmid pGL4.45 (luc2P/ISRE) or pGL4.32 (luc2P/NF-κB-RE) together with the indicated expression plasmids for 24 h. (A) Reporter activity driven by TRIF, RIG-I or MDA5 in the presence of increasing amounts of IL-17RD (IL-17RD; wedges), expressed as a percentage of the response obtained without IL-17RD. (B) ISRE reporter activity driven by increasing amounts of TRIF, RIG-I or MDA5 in the presence of empty vector (EV) or a fixed amount of IL-17RD (100 ng). (C-D) ISRE (C) and NF-κB (D) reporter activity driven by TBK1, MAVS or the constitutively active IRF3-5D mutant in the presence of increasing amounts of IL-17RD. Relative luciferase activity was measured as described in Materials and Methods (n=3). Data are mean ± SD. *p < 0.05, **p < 0.01, ***p < 0.001, ****p < 0.0001; # p < 0.05 versus the corresponding control.

