## Supplemental Figures 1-5 for "The orphan receptor IL-17RD is a negative regulator of RIG-I-like receptor-dependent antiviral innate immunity and restrains SARS-CoV-2-induced lung inflammation"

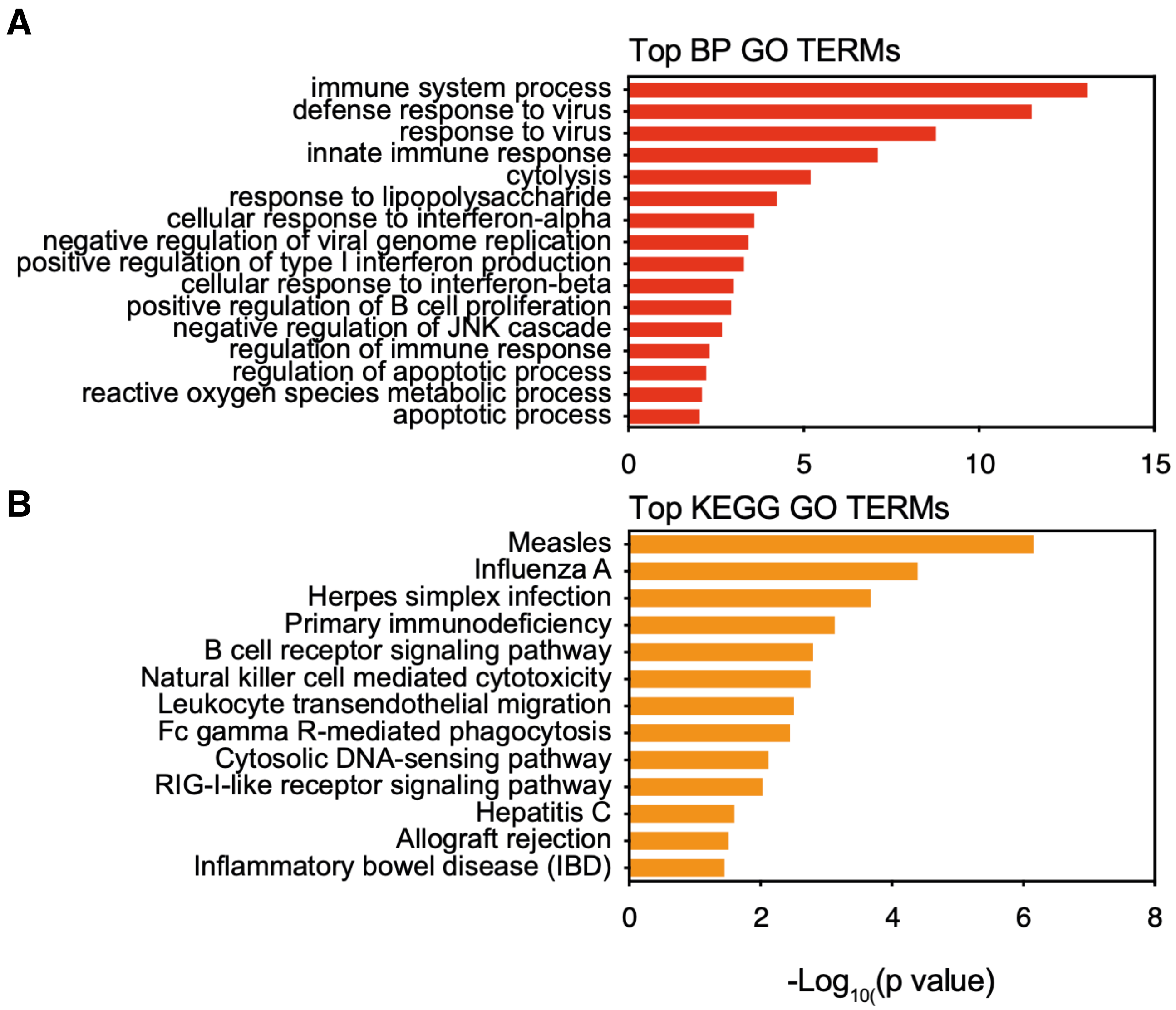


Supplemental Figure 1

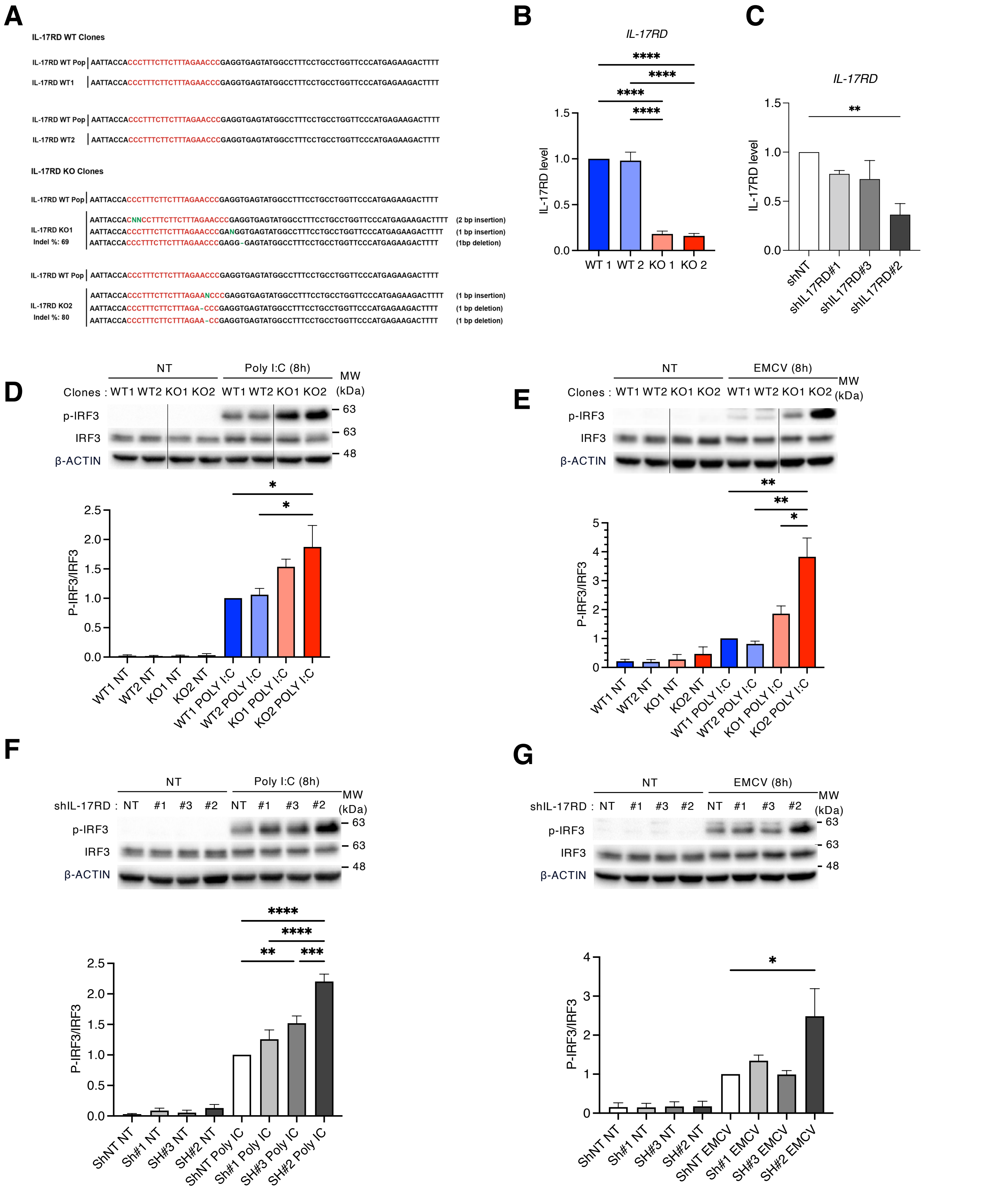


Supplemental Figure 2

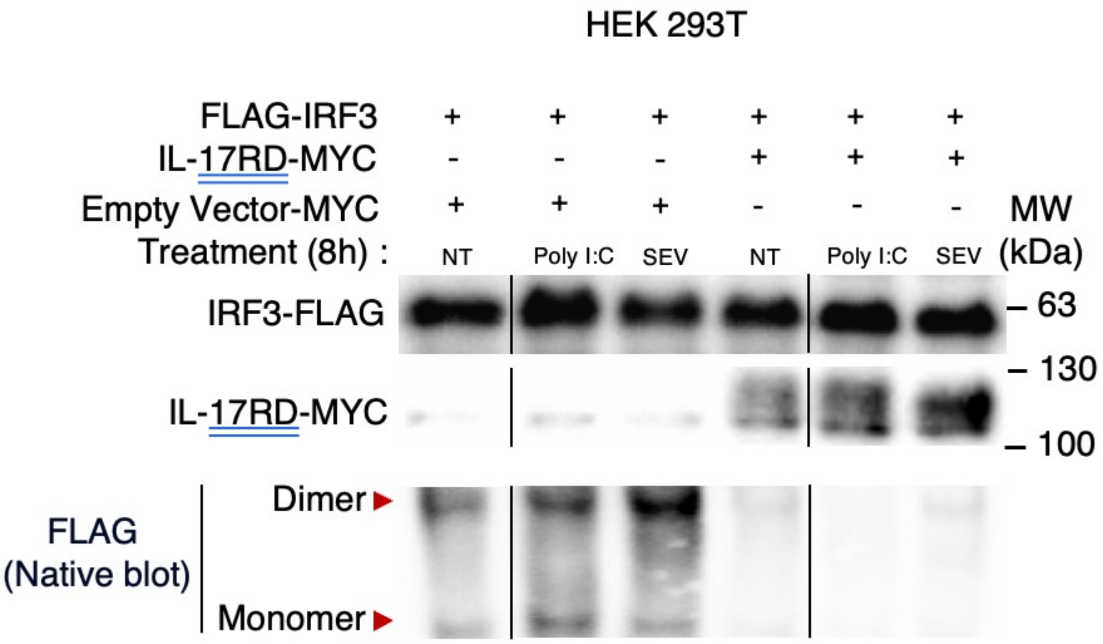


Supplemental Figure 3

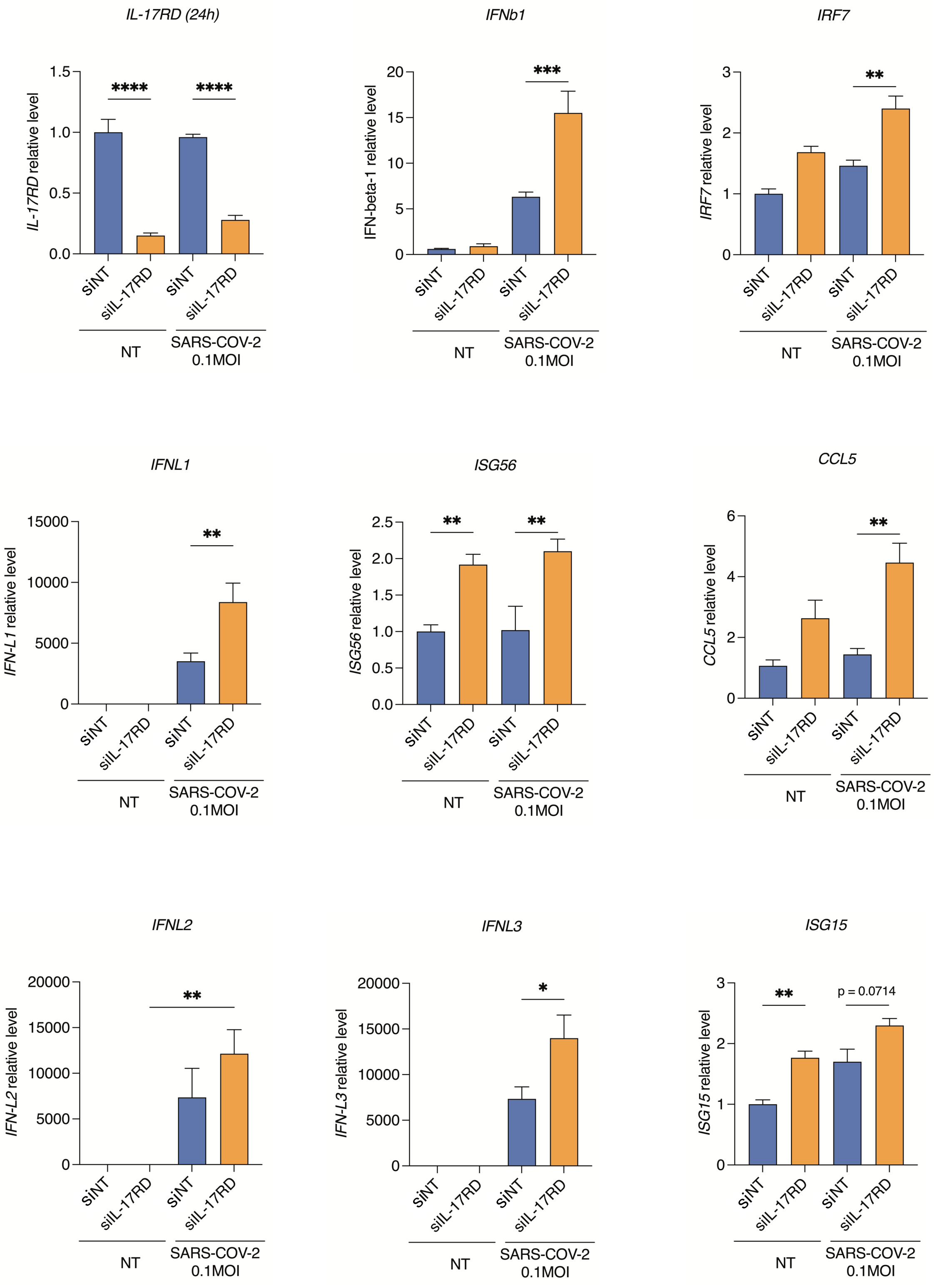


Supplemental Figure 4

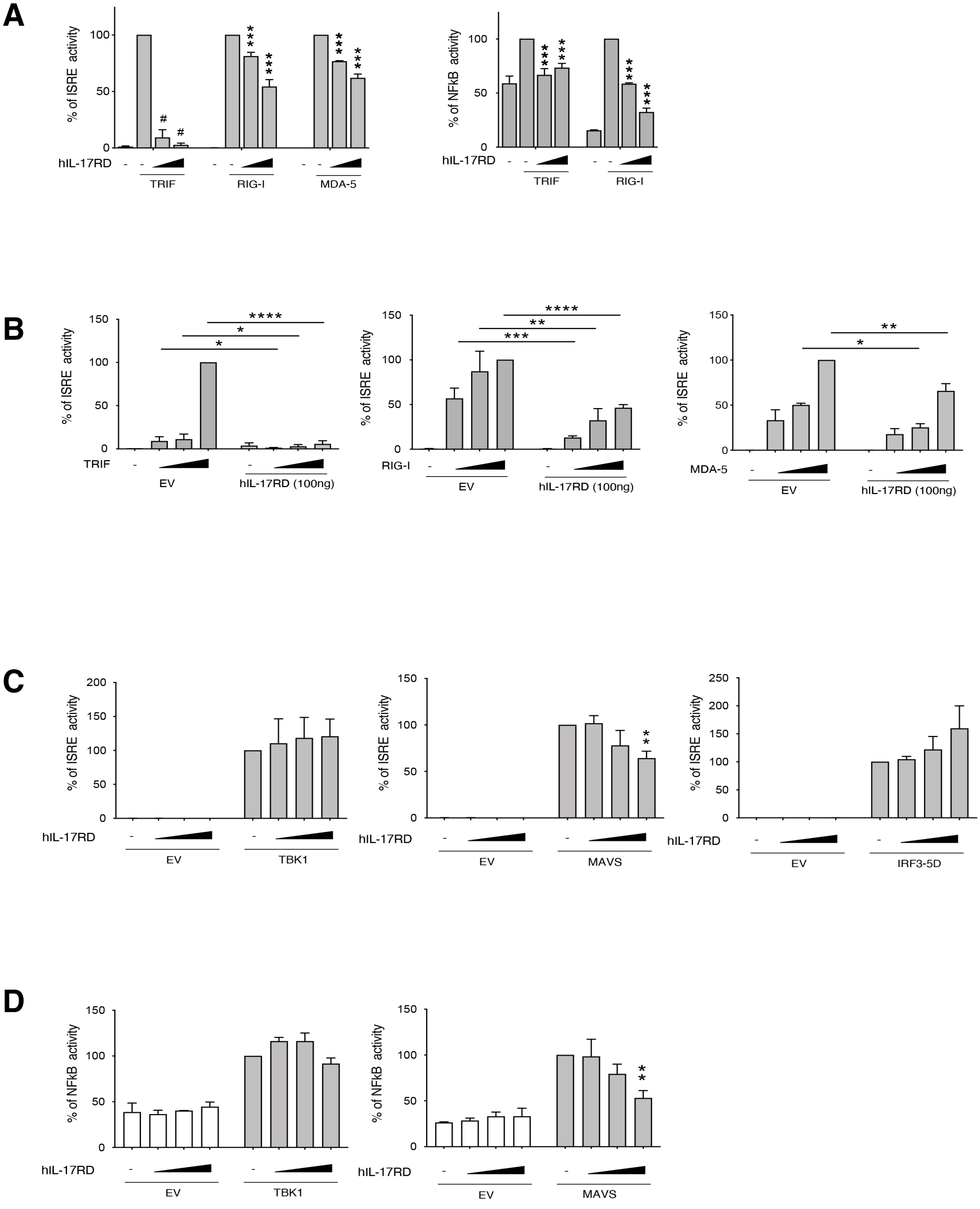


Supplemental Figure 5
